# Evidence for individual vocal recognition in a pair bonding poison frog? Insights from no-choice and two-choice paradigms

**DOI:** 10.1101/2023.09.15.558024

**Authors:** Molly E. Podraza, Jeanette B. Moss, Eva K. Fischer

## Abstract

Individually distinctive vocalizations are widespread in nature, although the ability of receivers to discriminate these signals has only been studied through limited taxonomic and social lenses. Here we ask whether anuran advertisement calls, typically studied for their role in territory defense and mate attraction, facilitate recognition and preferential association with partners in a pair bonding poison frog. Combining no- and two-stimulus choice playback experiments, we evaluated behavioral and physiological responses of females to male acoustic stimuli. Virgin females oriented to and approached speakers broadcasting male calls independent of caller identity, implying that females are generally attracted to male acoustic stimuli outside the context of a pair bond. When pair bonded females were presented with calls of a mate and a stranger, they showed a slight preference for calls of their mate. Moreover, behavioral responses varied with breeding status: females with eggs were faster to approach stimuli and spent more time in the mate arm than females that were pair bonded but did not currently have eggs. Our study suggests a potential role for individual vocal recognition in the formation and maintenance of pair bonds in a poison frog and raises new questions about how acoustic signals are perceived in the context of monogamy and biparental care.

## INTRODUCTION

Acoustic communication plays a prominent role in territory defense, mate choice, and social cohesion across animals (Simmons *et al*., 2003). Being both long-range and information-dense, acoustic signals confer distinct advantages over other types of signals and in many cases it is valuable for individuals to have their own ‘vocal signature’ that distinguishes them from others. Both the use of acoustic signals and individual vocal recognition have evolved independently multiple times across the animal kingdom, including in birds, mammals, fish, and amphibians (reviewed in Carlson *et al*., 2020; Chen & Wiens, 2020). Yet the importance of individual vocal recognition depends on a species’ social system, and the ability to produce and recognize distinct vocalizations should only evolve when more general recognition systems will not suffice (Carlson *et al*., 2020). Even within populations, individual vocal recognition has been shown to vary according to sex (Insley *et al*., 2003; Freeman & Ophir, 2021), age (Sieber, 1986; Balcombe, 1990; Leonard *et al*., 1997), and reproductive status (Pultorak *et al*., 2017; Shave & Waterman, 2017), and may be context-dependent. For instance, single male prairie voles are able to discriminate between male conspecifics but not between female conspecifics (Zheng *et al*., 2013), yet pair bonded males can discriminate conspecifics independent of their sex (Blocker & Ophir, 2015). Thus, evaluating how animals respond to acoustic stimuli and discriminate between them is a good way to test predictions about the selective context in which signals evolve.

One social context expected to strongly influence the evolution of individual recognition is pair bonding (Prior *et al*., 2022). Both the formation and maintenance of a pair bond depends critically on an individual’s ability to recognize and preferentially associate with a specific partner, which may be facilitated through one or more signal modalities (Prior *et al*., 2022). As pair bonding frequently co-occurs with biparental care, multi-modal signaling may also facilitate the coordination of complex care to ensure offspring survival (Boucaud *et al*., 2016, 2017; Moss *et al*., 2023). Taken together, the ability to distinguish and respond rapidly to partner vocalizations may contribute directly to fitness in monogamous and biparental systems, providing a selective context favoring the evolution of individual vocal recognition. Support for this idea can be found in monogamous birds, among which use of acoustic cues to recognize mates is widespread (e.g., Beer, 1971; Curé *et al*., 2011; Curé *et al*., 2016; Dentressangle *et al*., 2012; Lengange *et al*., 1999; Szipl *et al*., 2014; Wooller, 2010). In possibly the most well-studied example, zebra finches form remarkably enduring pair bonds, which are maintained *via* individuals’ strong preference for the songs of their mates (Miller, 1979; Hernandez *et al*., 2016) even in the absence of visual contact (Silcox & Evans, 1982) and when presented only with unlearned calls (D’Amelio *et al*., 2017). Discrimination of partner vocalizations is also observed in mammals (Snowdon & Cleveland, 1980; Snowdon *et al*., 1983), but evidence from other taxa remains limited despite multiple independent origins of monogamy and parental care (Emlen & Oring, 1977).

Anurans (frogs and toads) have long served as models for the study of acoustic communication. While the contexts in which frogs vocalize are varied and numerous, the extent of sensory reliance on acoustic signals (i.e., as opposed to other modes of communication, such as visual (Taylor *et al*., 2011)) and powers of individual discrimination remain poorly resolved. For example, only some species appear to discriminate between calls of neighbors and strangers (Lesbarrères & Lodé, 2002; Bee *et al*., 2016; Tumulty *et al*., 2022) – a pattern that has been linked to living in more complex social environments with strict territorial boundaries (Tumulty & Bee, 2021). Moreover, almost nothing is known from this group about individual vocal recognition in other social contexts, such as pair bonding.

The mimic poison frog, *Ranitomeya imitator*, is a monogamous species in which males and females form exclusive pair bonds lasting many months, defend shared territories, and cooperate over the care of eggs and tadpoles within those territories (Brown *et al*., 2008, 2010). Acoustic communication plays a key role in coordinating these behaviors, with males (the sole vocalizers in this species) calling to attract, court, and solicit trophic egg laying from females, who typically follow male partners at a close clip while they call (Brown *et al*., 2008; Tumulty *et al*., 2014). Recent work revealed that cooperation between pair bonded partners, specifically coordinated biparental egg-feeding, has contributed to the diversification of male signals (Moss *et al*., 2023). However, considerable overlap between call types and individual callers in the same study leaves open questions about whether receiver (female) recognition and/or preference has co-evolved alongside acoustic signals and to what extent multimodal cues (i.e., visual and/or olfactory) are necessary for individual-level recognition.

With this study, we test whether females of *R. imitator* (1) behaviorally respond to playback of male calls in general (i.e., when not pair bonded), and (2) discriminate and preferentially associate with calls of a mate over calls of a stranger. We combine single-stimulus (no-choice) and two-stimulus (two-choice) test paradigms to evaluate responses of virgin and pair bonded females to male acoustic stimuli. Choice tests are a standard method for evaluating female preference in anurans (Gerhardt, 1995; Bush *et al*., 2002) and other taxa (Wagner, 1998), and use of simple playback stimuli ensures subjects are responding to acoustic cues alone, as opposed to visual or olfactory cues (Carlson *et al*., 2020), the roles of which are unknown in our system. Our results provide a first glimpse into how vocalizations and use of individual vocal signatures may facilitate pair bond formation and maintenance in a biparental frog species.

## MATERIALS AND METHODS

### Subjects

All experimental frogs were sourced from a captive poison frog colony at the University of Illinois Urbana-Champaign. Founders of our colony descended from hybrids of five localities and were reared in admixed social groups. Although color morph variation has been suggested to influence mate choice in geographically isolated populations in the wild (Twomey *et al*., 2014, 2016), based on the breeding success of these laboratory-established pairs, we conclude that any effect of morph differences on preference formation and maintenance in our study is negligible. Adults were housed in opposite-sex pairs in 12x12x18” glass terraria (Exo Terra, Mansfield, MA, USA) containing a bioactive substrate layer (mix of soil, clay, orchid bark), sphagnum moss, live plants, dry film canisters for egg-laying, and water-filled film canisters for tadpole deposition. Froglets were group-housed (up to 10 frogs in plastic container with the same bioactive substrate and live plants) until 6–8 months of age, at which point they were transferred to adult terraria and housed in same-sex pairs or trios during the experiment. Animal rooms were temperature-controlled (21–23°C) on a 12L:12D cycle, and terraria humidity was maintained at >80%. Adults were fed wingless *Drosophila* fruit flies dusted with multivitamin and calcium supplements three times per week. Froglets were fed springtails and (after attaining sufficient body size) fruit flies three times per week.

### Acoustic Stimuli

The advertisement calls of 12 adult, breeding males were used as stimuli for choice tests. Relative to other call types in the repertoire of *R. imitator* (e.g., courtship or egg feeding calls), advertisement calls contain significant identity information (Moss *et al*., 2023) but show weak or no differences between color morphs (Twomey *et al*., 2015). Therefore, we expected advertisement calls to be a reliable signal for distinguishing individuals. We followed our previously established protocol for recording frogs in the laboratory (Moss *et al*., 2023). Briefly, subjects were recorded in their home tanks during daytime hours using a digital audio recorder (H4n Pro, Zoom, Tokyo, Japan) and shotgun microphone (K6/ME66, Sennheiser, Wennebostel, Germany) positioned directly above acoustically transparent screen mesh lids (<0.5 m). To generate playback files, we used the open-source audio editing software Audacity v. 3.0.4 (Audacity Team, 2021). Recordings were filtered to remove background noise using the noise reduction, high-pass filter, and low-pass filter tools. We then selected ten high-quality calls per male and applied a standardized call rate of ten seconds to all playback files (Fig. 1A) to control for variation in signal perception introduced by call rate or sampling variance. Given that most spectral and temporal properties of advertisement calls except for inter-call interval are shown to have high potential for individual coding in *R. imitator* (Moss *et al*., 2023), this protocol captures the range of meaningful within-individual variation in calls. All stimulus files were matched for peak amplitude prior to playback.

**Figure 1:**
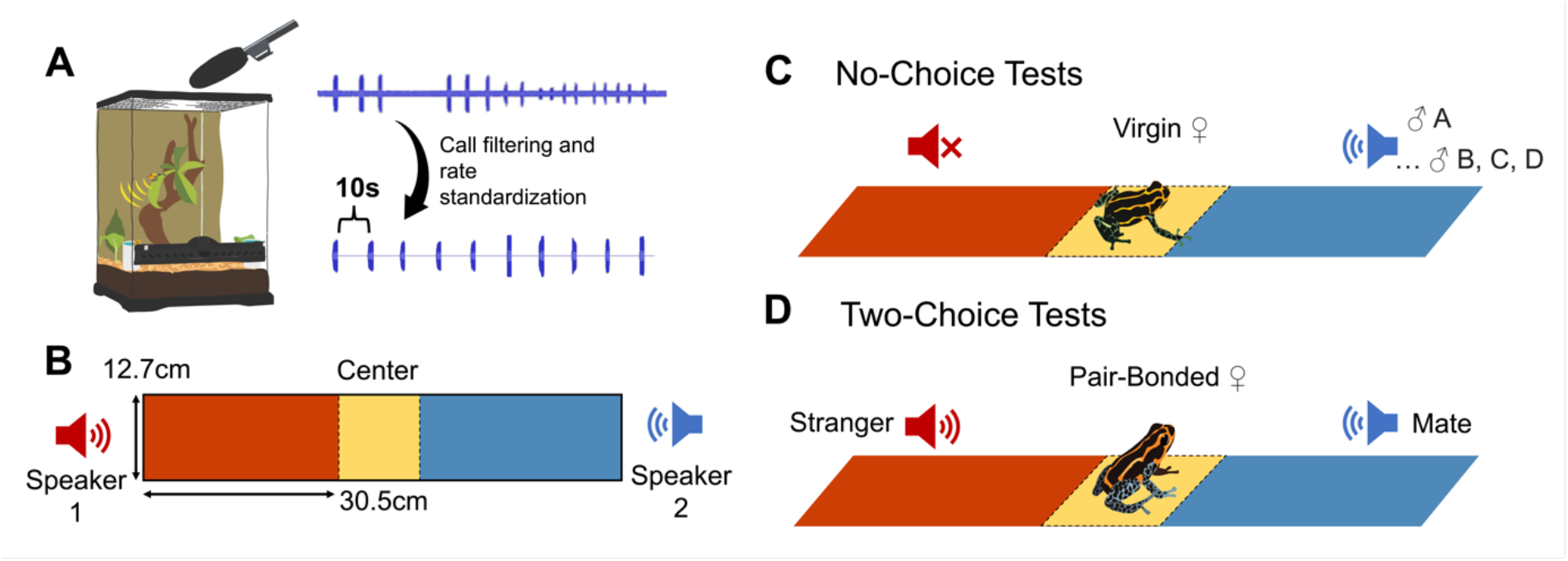
Experimental overview of choice tests. (A) Acoustic stimuli to be used for playback were generated from advertisement calls of adult male *Ranitomeya imitator*, which were recorded in the laboratory and filtered to remove background noise and standardize call rates. (B) Trials were conducted in a two-arm maze inside of a sound isolation booth, with subjects released in the center of the arena. (C) No-choice (single-stimulus) tests were conducted by broadcasting male acoustic stimuli from one (randomized) side of the arena. Subjects (virgin females) were presented with five male acoustic stimuli in a randomized order. (D) Two-choice (two-stimulus) tests were conducted by broadcasting competing acoustic stimuli from opposite (randomized) sides of the arena. Subjects (pair bonded females) were presented with acoustic stimuli of their mate and an unfamiliar male.

### Experimental Overview

#### Phonotaxis apparatus and procedures

All trials were conducted inside a sound isolation booth (SE 2000, WhisperRoom, Inc., Knoxville, TN, USA) furnished with 2” of acoustical foam to minimize ambient noise. The apparatus used for the choice tests was a long corridor measuring 29” long, 5” wide, and 7” tall with a central starting point, generating two equal-length arms (12” long) running in opposite directions (Fig. 1B). A wooden frame provided the main scaffold for the structure while acoustically transparent black mesh enclosed the walls. The floor was lined with the same substrate mixture and moss used in the frogs’ home terraria and was misted in between each trial to maintain consistent humidity levels. To facilitate video recordings, the top of the arena was cut from plexiglass and the interior was softly illuminated with LED tape.

Acoustic stimuli were broadcast from speakers (Mod1 Orb speaker, Orb Audio, New York, NY, USA) located just outside the end of the choice arms. Stimulus files were stored on individual USB flash drives such that at the end of each ten-call sequence, playback restarted from the beginning and looped for the duration of the trial. Calls were broadcast through amplifiers (MAMP1, MouKey, Solihull, UK) at 60–70dB, which approximates the sound pressure level of calling males measured from a distance of 12”. At the start of each trial, the focal female was placed under a clear deli cup in the center of the arena and playback was initiated from one or both speakers. For trials that utilized both speakers, competing stimuli were offset by five seconds to minimize interference. After an acclimation period, we used a pulley system – a string threaded through the plexiglass and operated by the observer concealed behind a privacy blind – to lift the cup, marking the start of the choice test. While tests of phonotactic behavior in anurans have traditionally involved explosive breeding species that orient and move rapidly to the first attractive sounds they hear (Gerhardt, 1995; Wells, 2010), the breeding biology of *R. imitator* (a pair bonding species with prolonged reproductive and courtship periods) is unlikely to have selected for such vigorous responses. To account for this, subjects were afforded ten minutes to move throughout the arena and their behavior quantified throughout, rather than as a single choice response. All trials were recorded using remote-controlled video cameras (Brave 7, AKASO, Frederick, MD, USA).

#### Experiment 1: No-Choice Tests

To evaluate female responses to male acoustic stimuli independent of pair bond status, we exposed virgin females (all 8–9 months old at the time of testing) to a single acoustic stimulus in a no-choice test. Subjects (N=9) were chosen from among female F1 offspring of colony founders. These individuals were reared in the lab without access to deposition pools and were never housed with an adult male, and thus we could be confident that attraction to acoustic stimuli was unaffected by a prior pair bond. Nevertheless, acoustic stimuli may vary in their attractiveness to females due to qualitative differences among males. To account for this possibility, we sequentially tested each subject’s response to each of five acoustic stimuli, each from a different, unfamiliar male (Fig. 1C).

We assayed each female over two consecutive days during the experimental period (June–July 2022) in alternating sets of two or three tests spanning 30–45 minutes. Due to attrition, only seven of nine subjects completed all five trials, while two completed only two or three trials. A single speaker was used for the stimulus and its placement was randomized between sides for each trial, such that the stimulus was broadcast from the left or right side in equal numbers. Tests were conducted in a randomized order between 10:00 and 13:00. Each test included a 5-minute acclimation period immediately prior to the 10-minute trial.

#### Experiment 2: Two-Choice Tests

To test whether pair bonded females preferentially associate with mate calls, we exposed adult females to competing acoustic stimuli, one of their mate and one of a stranger, in a two-choice test (Fig. 1D). Trials were conducted over two time blocks, nine weeks apart. Subjects were chosen from proven breeding pairs, defined as pairs that had successfully cared for at least one clutch (i.e., reared eggs to hatching and transported tadpoles). Three proven pairs could not be included in Block 2 owing to mortality of one or both partners. These were replaced with new breeding pairs who, in turn, were tested only in Block 2. In sum, N=9 subjects experienced between 2–4 tests (total N=28 trials). Breeding status at the time of testing (i.e., whether the pair currently had eggs or not) was recorded throughout.

For each block (February–April 2022 and May–June 2022), subjects experienced a series of choice tests. Tests were conducted in sets of three in a randomized order between 10:00 and 17:00. One of the tests involved white noise broadcast from one or both speakers; however, due to imperfect replication across blocks these results are not shown. We report here only on two tests that were performed with perfect replication: (1) A mate versus stranger test, in which calls of the mate were broadcast from the left side of the arena and calls of a stranger were broadcast from the right side; and (2) A stranger versus mate test, in which the same stimuli were used as in step 1 but speaker positions were reversed. This procedure was repeated in the second experimental period but with a different stranger stimulus. Thus, our design accounted for potential side preference as well as stranger identity preference. Each test included a 10-minute acclimation period in addition to the 10-minute recorded trial. The reason for this extended acclimation time was to account for the delayed exit from center by pair bonded females relative to virgins in our preliminary trials.

### Behavioral Analysis

We analyzed video recordings of trials using the behavioral analysis software BORIS (Friard & Gamba, 2016). Target behaviors were coded to reflect time spent (in seconds) in each arm of the phonotaxis arena (Fig. 1B). Entrance into a region was defined as the entire body of the subject crossing a superimposed threshold. Scoring was conducted blind with respect to treatment group and stimulus side.

We imported ethogram outputs from BORIS into R Studio v. 3.386 (R Core Development Team, 2020) for statistical analysis. For each trial, we quantified the total duration of time frogs spent in each arm of the phonotaxis arena. For the no-choice tests, the two arms represented “Towards Stimulus” and “Away from Stimulus”. For the two-choice tests, the arms represented “Toward Mate” and “Toward Stranger”. To assess whether subjects showed an initial preference between arms and how quickly this judgement was made, we also extracted the first movement out of the center and, for no-choice trials, the first movement towards the stimulus, for each trial. We compared time spent between arms and the latency to first move into either arm by fitting mixed effects models in lme4 (Bates *et al*., 2015), where breeding status was included as a covariate (for the two-choice tests) and female ID, speaker position, and block (for the two-choice tests) were included as random effects. The number of first movements in either direction for each block was compared against chance levels using a Chi-square test. To understand how phonotaxis behavior varied as a factor of male ID in the no-choice tests, we fitted a mixed effects model as above and significance was evaluated *via* ANOVA followed by post-hoc testing with the package emmeans (Lenth *et al*., 2022).

To account for changes in space use over time (i.e., directional movement towards or away from a given stimulus), we recalculated duration of time spent in each region of the arena for the first, middle, and last 200 seconds of a trial and modeled time in each region as a function of location, time, and their interaction with female ID, speaker position, and block (for two-choice tests) specified as random effects. We again evaluated significance *via* ANOVA followed by post-hoc testing with the package emmeans.

## RESULTS

### Experiment 1: No-choice tests

Virgin females spent more time in the arm with the stimulus than without (Fig 2A; t_3,37_ = 7.529, *p* = 0.008) and there was no significant side bias. Identity of the male caller had no significant impact on either the latency of females to approach the stimulus (F_3,31_ = 1.10, *p* = 0.375) or on the time females spent in the arm broadcasting the stimulus (F_3,37_ = 1.183, *p* = 0.345). Subjects also did not significantly alter their space use over time (Fig. 2B; Tukey HSD *p* > 0.05). Rather, frogs consistently spent more time toward the stimulus than away from it, and this difference attained significance by the final 200 seconds of the test (Fig. 2B; Tukey HSD t = 2.424, *p* = 0.044). While in the majority of trials (N=29) the first movement was directed towards the stimulus as opposed to away from it (N=13; χ^2^ = 3.270, *p* = 0.071), approach was slow regardless of initial orientation (F_2,35_ = 0.009, *p* = 0.925; mean latency to exit center = 144 s, range = 2–548 s). In three of 40 trials, the focal female did not move from center for the full ten minutes. Random effects (female ID and speaker position) together accounted for <10% of residual variance in the overall distribution of time spent in either arm (<0.005) or in latency to exit center (0.086).

**Figure 2:**
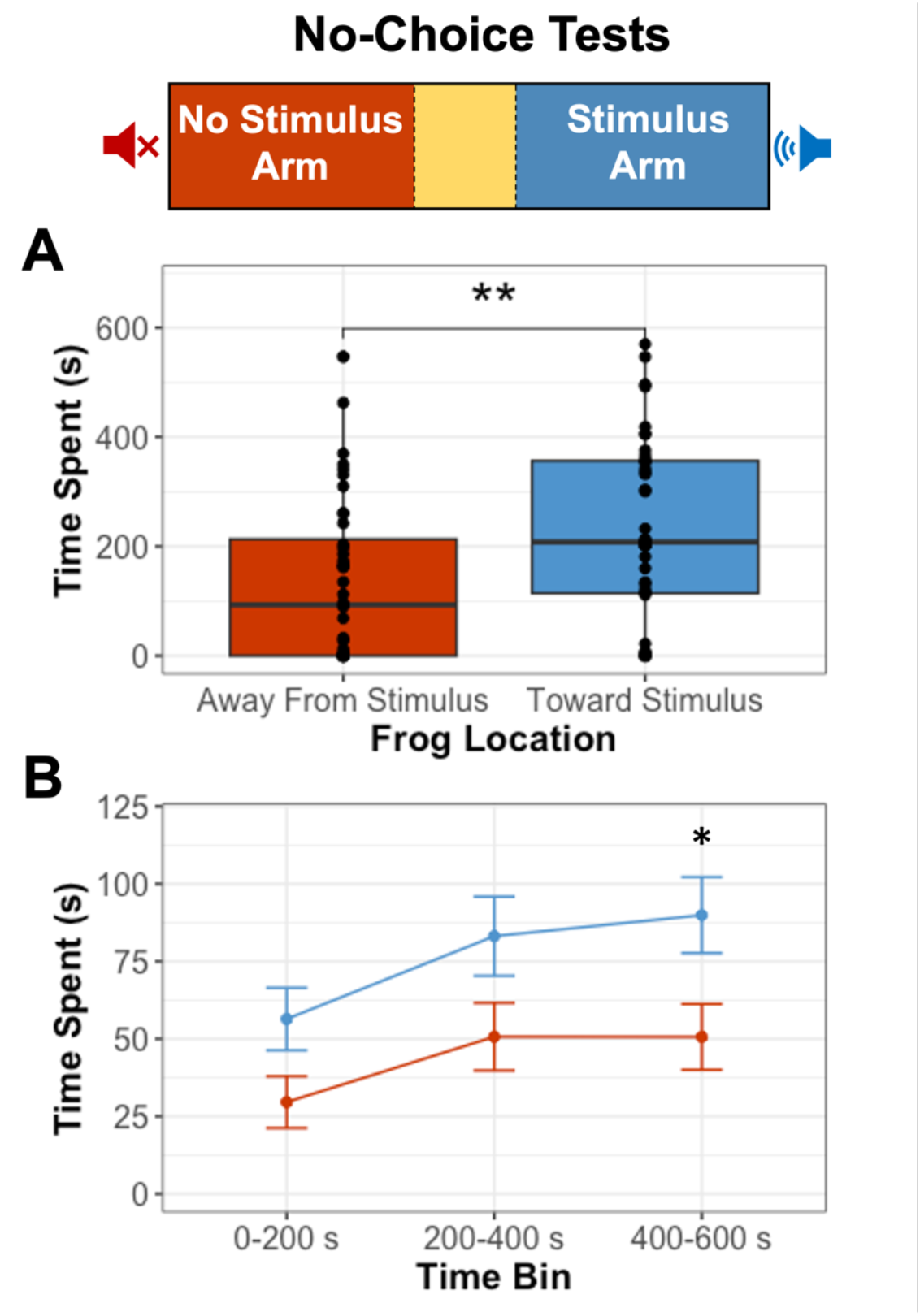
Summarized results from the no-choice (single-stimulus) tests of virgin female *Ranitomeya imitator*. (A) Cumulative time spent in the arm without an acoustic stimulus (red) compared to the arm with an acoustic stimulus (blue), in seconds. (B) Changes in frog location over time, depicted as the cumulative time spent in the no stimulus (red) versus stimulus (blue) arm in the first, middle, and last 200 seconds of the trial. Means and standard errors are depicted for each time bin, and asterisks denote significant differences between arms within a given time bin.

### Experiment 2: Two-choice tests

There was a borderline significant trend of pair bonded females spending more time in the arm broadcasting acoustic stimuli of their mate than that of a stranger (Fig. 3A; F_4,24_ = 3.687, *p* = 0. 060). Closer inspection of the time course of female behavior revealed that subjects significantly favored the mate arm for the first 400 seconds of the trial (Tukey HSD: 0–200 s: t = 2.999, *p* = 0.010; 200–400 s: t = 2.762, *p* = 0.019) but increased time spent in the stranger arm in the final 200 seconds (t = 3.205, *p* = 0.005), diminishing overall differences between stimuli (Fig. 3B). Initial orientation was more frequently directed into the mate arm (N=12) than the stranger arm (N=7), albeit not significantly so (χ^2^ = 1.315, *p* = 0.251). Initial orientation had no bearing on the latency to first movement (F_4,15_ = 1.945, *p =* 0.185); however, subjects that initially oriented towards the mate arm spent cumulatively more time there (F_3,16_ = 1.945, *p* = 0.005) as did subjects with shorter latencies to exit the center (F_3,16_ = 16.425, *p* = 0.0008).

**Figure 3:**
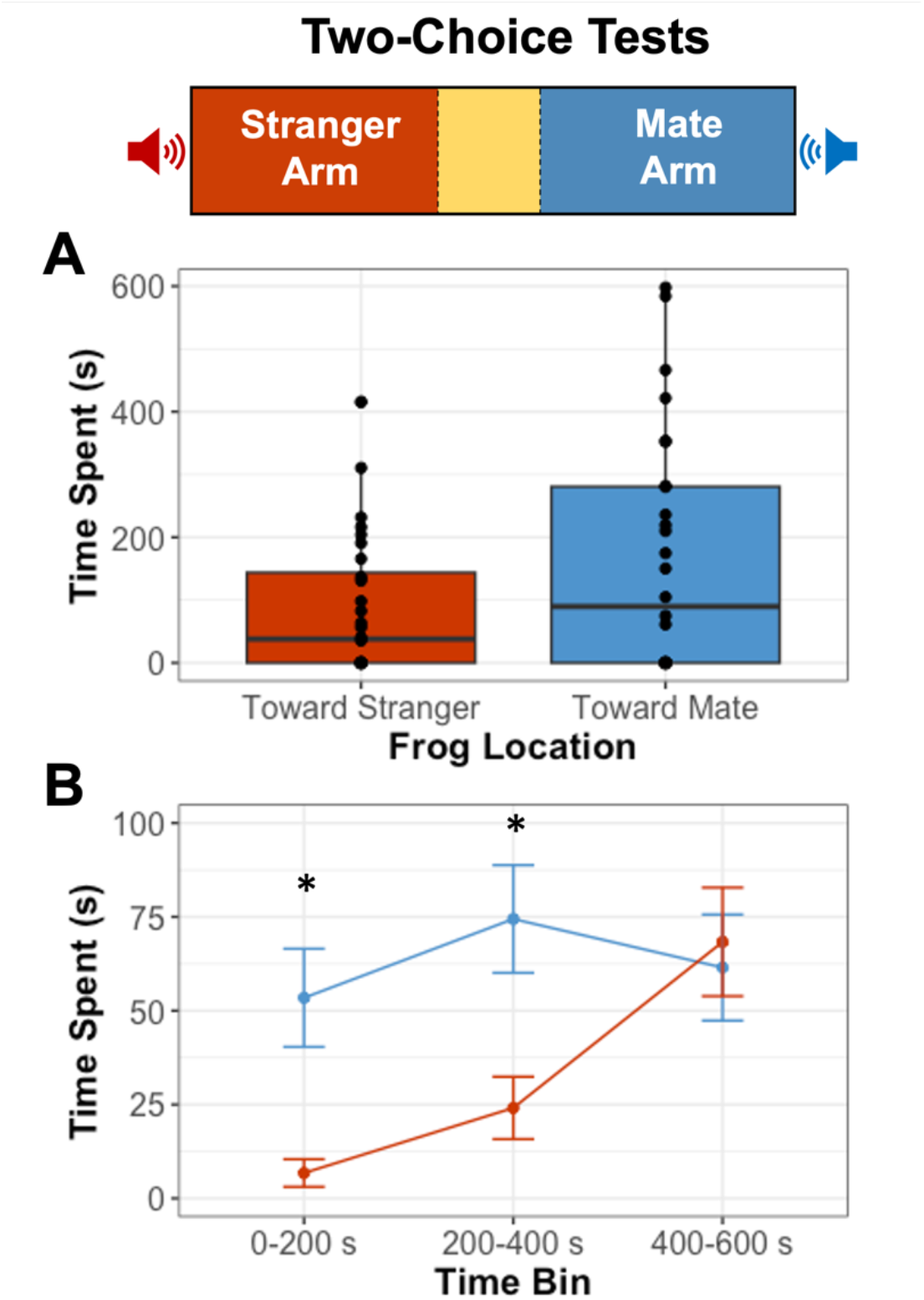
Summarized results from the two-choice (two-stimulus) tests of pair bonded female *Ranitomeya imitator*. (A) Cumulative time spent in the arm broadcasting a stranger stimulus (red) compared to the arm broadcasting a mate stimulus (blue), in seconds. (B) Changes in frog location over time, depicted as the cumulative time spent in the stranger (red) versus the mate (blue) arm in the first, middle, and last 200 seconds of the trial. Means and standard errors are depicted for each time bin.

**Figure 3:**
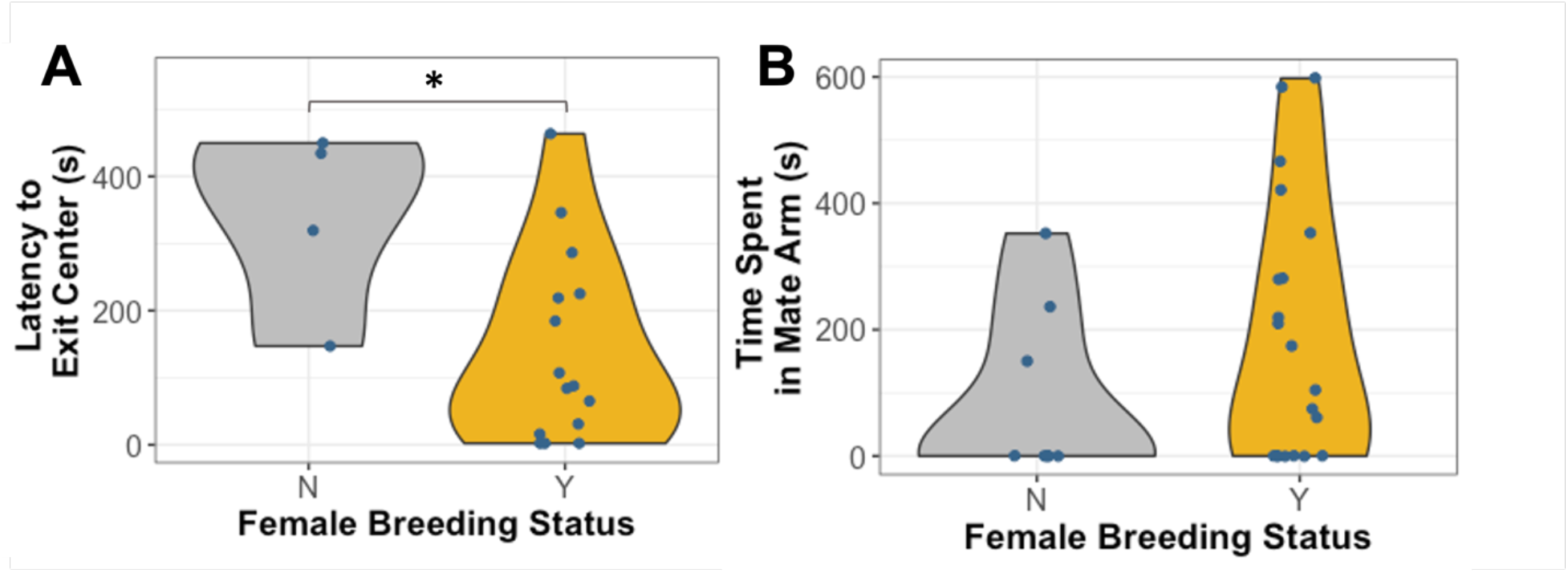
Effect of breeding status (N = female does not currently have eggs; Y = female currently has eggs) on behavior of pair bonded female *R. imitator* in phonotaxis arena. (A) Latency to exit the center of the arena and enter either arm, in seconds; (B) Cumulative time spent in the arm broadcasting a mate stimulus, in seconds.

Females that had eggs at the time of testing were significantly faster in exiting the center (F4,15 = 6.80, *p* = 0.020; Fig. 3A) and spent marginally more time in the mate arm (F4,24 = 3.454, *p* = 0.069; Fig. 3B) than females who were pair bonded but did not have eggs or tadpoles at the time of testing. Random effects (female ID, block, and speaker position) together accounted for < 12% of residual variance in the overall distribution of time spent in either arm (< 0.005) or latency to exit center (0.112). We note that for nine of 28 trials in Experiment 2, the focal female did not move from center for the full ten minutes, which is a significantly higher proportion than in Experiment 1 (Fisher odds ratio = 5.682, *p* = 0.011). However, this behavior could not be attributed to specific subjects or breeding status.

## DISCUSSION

In this study, we performed playback experiments in the laboratory to evaluate the scope for vocal recognition at the individual level in the pair bonding poison frog, *Ranitomeya imitator*. Through a series of no-choice experiments using virgin females as test subjects, we first demonstrated that acoustic signals elicit behavioral responses from inexperienced females and that male identity does not contribute significantly to variation in preference for unfamiliar individuals. Following demonstration of a general preference for conspecific males, we subjected pair bonded females to two-choice tests involving acoustic stimuli of their pair bonded mate versus a stranger. With this design, subjects were more reticent to explore the arena, and many appeared to sample both stimuli rather than preferentially associating with one. Nonetheless, we found a trend wherein females spent more time near the speaker broadcasting playback of their mate than that of a stranger, especially early on in the trial. Ours is the first study to show that females may be able to distinguish between males based on calls alone, suggesting a role for individual recognition in partner preference across poison frogs.

The primary motivating question of our study was whether pair bonded females would discriminate and preferentially associate with calls of their mate – a phenomenon that has been documented extensively in monogamous birds and some mammals (Beer, 1971; Miller, 1979; Snowdon & Cleveland, 1980; Snowdon *et al*., 1983; Lengange *et al*., 1999; Wooller, 2010; Curé *et al*., 2011, 2016; Dentressangle *et al*., 2012; Szipl *et al*., 2014; Hernandez *et al*., 2016). The first step in testing this was to determine whether female *R. imitator* are attracted to male acoustic stimuli in general, independent of pair bonding. To our knowledge, only one experiment to date has examined behavioral responses of *R. imitator* to acoustic playback, and this study focused on interactions between territorial males in the field (Mayer *et al*., 2014). Here, we show that virgin female *R. imitator* preferentially associate with male acoustic stimuli when the alternative is no sound. Behavioral responses did not vary significantly among subjects or between the five distinct male stimuli presented, suggesting that any advertisement call is sufficient to elicit interest from inexperienced females. Use of subtle variation in male advertisement calls to assess mate quality has been demonstrated experimentally in related poison frogs (Forsman & Hagman, 2006; Dreher & Pröhl, 2014; Pettitt *et al*., 2019; Peignier *et al*., 2022), although these assessments may have little bearing on mate choice *in situ* (i.e., relative to practical factors, such as proximity; Meuche *et al*., 2013). Given the prolonged courtship of *R. imitator*, initial attraction to acoustic stimuli is only the first step in a more thorough evaluation of mate quality, which may involve visual and olfactory cues of which more experienced females may be more discerning. Future studies will be necessary to determine the role of acoustic signal variation in initial mate attraction in *R. imitator* and the influence of experience on choice.

In our two-choice paradigm, we found that pair bonded females spent more time associating with the mate stimulus than the stranger stimulus, that this was especially true immediately after stimulus presentation, and that this behavior was correlated with initial orientation towards the mate stimulus (Fig. 3A). A similar, non-significant trend was recently found in a study of a promiscuous dendrobatid (*Allobates femoralis*): that females tend to prefer familiar partners over novel partners as mates (Peignier *et al*., 2022). The authors suggest this result may be due to captive housing conditions and/or knowledge of males’ territorial status. Alternatively, females may be picking up on acoustic indicators of prior paternal experience, as has been shown in *Dendrobates leucomelas, Epipedobates tricolor*, and *Anomaloglossus beebei* (Forsman & Hagman, 2006; Pettitt *et al*., 2019). Compared to non-breeding females, females in our study that currently had eggs were quicker to exit the center and showed stronger preference for their mate. This raises the intriguing possibility that physiological changes associated with reproduction (i.e., hormonal changes) could enhance sensory perception of and/or motivation to respond to mate stimuli (e.g., Dey *et al*., 2015). Even so, considerable variation within and between individual trials leads us to conclude that females probably rely on multimodal cues (e.g., visual and olfactory) for accurate identity determination in most circumstances. Such would be consistent with the mating and social system of *R. imitator*, as males and females cohabit small territories that are typically non-overlapping with other pairs (Brown *et al*., 2008, 2010) and the need to distinguish partners based on calls alone may arise infrequently.

An important caveat to the above findings is that in almost a third of trials, pair bonded females did not move from the central starting point. When subjects did explore the arena, they spent less time outside of center than females in no-choice trials, and the difference in time spent between arms was not statistically significant (Fig. 3A). Reduced effect sizes in two-stimulus tests relative to single-stimulus tests is not without precedent in studies involving anurans (Tanner *et al*., 2017) and may result from females ‘‘sampling” both sides and obscuring choice (Bush *et al*., 2002). In line with this, inspecting the time course of frog locations in the arena revealed investigation of the stranger stimulus increased only at the end of trials (Fig. 3B), suggesting that a female’s motivation to approach a stranger may stem from their failure to locate their mate or from a place of territory defense. For instance, monogamous California mice that use ultrasonic vocalizations for territory defense do not preferentially approach vocalizations of their partners but do approach vocalizations of a stranger (Pultorak *et al*., 2017).

In conclusion, our study shows that females of a pair bonding poison frog are behaviorally responsive to male acoustic signals and, for the first time in an anuran, evaluates individual vocal recognition in the context of monogamy. Pair bonding is predicted to strongly influence the evolution of sensory perception and communication (Prior *et al*., 2022), although empirical tests of this prediction have been rare outside of birds and mammals. Having previously shown that advertisement calls of *R. imitator* are reliable signals of male identity (Moss *et al*., 2023), weak female preference between male vocalizations in the present study suggests that additional signal modalities and context cues are vital to the maintenance of pair bonds in this system. At the same time, behavioral motivation must be interpreted through the lens of species biology, and ultimately the ambiguity of our results highlights limitations of translating standard behavioral assays across taxa. Given *R. imitator*’s prolonged reproduction and courtship, variation in female assessments of male calls may be more evident at the hormonal or neural, as opposed to behavioral level. Future investigations should explore the use of alternative assays for uncovering the role of acoustic and other modes of communication in maintaining pair bond structures.

## ACKOWLEDGEMENTS

We thank Mark Bee for helpful advice on phonotaxis and playback equipment during early project design stages and the Machine Shop at University of Illinois Urbana-Champaign, in particular Scott Baker, for constructing the experimental apparatus.

## DATA AVALIABILITY

All raw data and R scripts will be uploaded to Data Dryad and made publicly available upon article acceptance.

